# WayFindR: Investigating Feedback in Biological Pathways

**DOI:** 10.64898/2026.03.27.714788

**Authors:** Polina Bombina, Reginald L. McGee, Jake Reed, Zachary B. Abrams, Lynne V. Abruzzo, Kevin R. Coombes

**Affiliations:** Department of Biostatistics, Data Science, and Epidemiology, Georgia Cancer Center at Augusta University, Augusta, GA 30912, USA; Department of Mathematics and Statistics, Haverford College, Haverford, PA 19041, USA; Department of Oncological Sciences, Huntsman Cancer Institute, Salt Lake City, UT 84112, USA; Institute for Informatics, Data Science & Biostatistics, Washington University, St. Louis, MO 63110, USA; Department of Pathology, Medical University of South Carolina, Charleston, SC 29425, USA

**Keywords:** Wikipathways, KEGG, homeostasis, negative feedback, feedback loops

## Abstract

Understanding biological pathways requires more than static diagrams. We present `WayFindR`, an R package that converts pathway data from WikiPathways and KEGG into graph structures using `igraph`, enabling computational analysis of regulatory features such as negative feedback loops. Rooted in control theory, negative feedback is essential for system stability, yet it is often underrepresented in curated pathway data.

In this study, we systematically analyzed pathway information from both databases across multiple species and found that feedback loops—particularly negative ones—are rarely captured. This gap likely reflects both biological and technical challenges. Biologically, feedback mechanisms are inherently complex and often remain uncharted due to limited experimental focus. Technically, pathway databases frequently lack standardized annotations and complete representations of regulatory interactions, especially inhibitory edges that are crucial for identifying feedback.

These observations underscore the need for improved data curation and consistent annotation practices to enhance our understanding of regulatory dynamics. By bridging the gap between static pathway diagrams and dynamic systems-level insights, `WayFindR` enables reproducible and scalable investigation of feedback regulation in cellular networks.

The `WayFindR` R package can be downloaded from the Comprehensive R Archive Network (CRAN) (https://cran.r-project.org/web/packages/WayFindR/index.html). The processed data along with code for download can be accessed via the GitLab repository (https://gitlab.com/krcoombes/wayfindr).

## 1. Introduction

Homeostasis, the self-regulating process by which an organism maintains internal stability while adjusting to changing external conditions, was defined by Walter Cannon in 1926 [9]. By now, it has become a central concept of biology and physiology [3]. Early on, Cannon emphasized that homeostasis is not something set and immobile. Rather, it is a dynamic condition that may vary, yet stays relatively constant [5]. Cannon’s introduction of homeostasis shifted the focus of physiology from the sense of a “fixed value” to a system of countervailing forces keeping the values within a narrow range [9].

Integral to understanding how the body maintains homeostasis is the notion of feedback [9]. As Jamie Davies wrote [11], “Beyond Earth, life without DNA is just about thinkable (one can imagine alternative strategies for storing information). Life without feedback loops, though? I have never met any biologist who can imagine that.” However, as George Billman wrote [3], “It is also important to note that homeostatic regulation is not merely the product of a single negative feedback cycle but reflects the complex interaction of multiple feedback systems that can be modified by higher control centers. This hierarchical control and feedback redundancy results in a finer level of control and a greater flexibility that enables the organism to adapt to changing environmental conditions.”

Feedback can be either positive or negative. Positive feedback occurs when the activation or accumulation of one component leads to the activation of another, amplifying the effect. If the only feedback in a system is positive, then the system tries to spiral off to infinity, and may eventually find itself limited by outside factors such as a lack of resources. For instance, a cell can only continue transcribing DNA to RNA if enough nucleotides are available; it can only continue translating RNA into protein if it has an adequate supply of transfer RNAs. In contrast, negative feedback occurs when the activation or accumulation of one component results in the deactivation or inhibition of another, thereby stabilizing the system [8]. It is these inhibitory effects that are essential to homeostasis, by attempting to restore a stable state. The mathematical discipline of control theory highlights the idea that negative feedback is crucial for the effective regulation and control of dynamical systems, ensuring that deviations from desired states are minimized and the system returns to equilibrium [14].

When talking about feedback, we are usually referring to feedback *loops*. That is, situations where downstream elements in a process can interact with upstream elements to enhance or inhibit their effects. The idea of feedback loops has a long history that extends across many branches of science and engineering. From James Watt’s addition of a “governor” to his steam engine [38] to the U.S. Navy engineers in the 1920’s and 1930’s working on gunnery control systems [29] to the thermostat controlling the temperature in your house, feedback loops have become ubiquitous. A search of Google Scholar for “feedback loops” returns 3,500,000 results. In the biological sciences, the same search in PubMed returns a steadily growing list of 22,678 results (**Figure 1**).

**Fig. 1.**
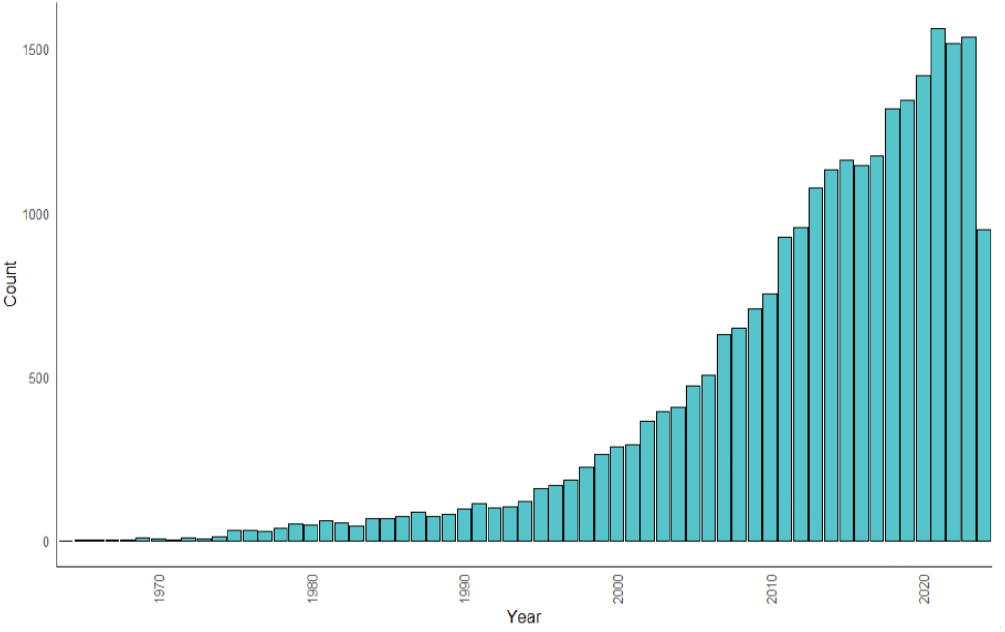
Trend in the number of publications related to feedback loops in PubMed by year.

The goal of this paper is to test the hypothesis, implicit in the quote from Jamie Davies above, that negative feedback loops should be common in intracellular biological pathways. One caveat, however, that needs to be kept in mind can be illustrated by the cell cycle. We know that the transition from one phase to another involves first stimulating the proteins required for the next phase and then inhibiting the proteins that were only needed for the previous phase. That part involves canonical feedback loops. But the cycle is also subject to oversight by “checkpoint” proteins, which monitor items like cell size, DNA replication, and DNA damage, and can stop the cycle if they detect problems [1]. An immediate “stop” signal is inhibitory, but need not be involved in a loop that provides feedback to earlier parts of the pathway. So, to test our hypothesis, we will need to track both loops and inhibitory interactions as potentially independent control features.

Much of our knowledge of biological pathways has been locked away in drawings and images in the biological literature, accessible only as files formatted using Joint Photographic Experts Group (JPEG) or Scalable Vector Graphics (SVG) formats that are not amenable to computational study [36]. The initiation of the Kyoto Encyclopedia of Genes and Genomes (KEGG) pathway database in 1995 began the process of making it possible to compute on pathways [22]. KEGG’s primary focus from the beginning has been on metabolic pathways, but they have gradually added regulatory pathways. The launch of the Reactome database in 2004 brought additional tools and a stronger focus on the ability to compute on biological pathways [21]. The fundamental unit of both the KEGG and the Reactome databases, however, is a *reaction*. This design decision necessarily emphasizes metabolites and small molecules over genes and proteins. By contrast, WikiPathways, introduced in 2008, uses a database model in which genes and proteins are given importance equal to metabolites or small molecules [36]. In this paper, we will analyze pathways from both WikiPathways and KEGG, both of which use a variant of the eXtensible Markup Language (XML) to define file formats making it possible to transport and share pathway diagrams.

Our interpretation of “computing on biological pathways” has a somewhat different slant from most previous efforts. We are not trying to assemble pathways from previous knowledge or the literature or studies of gene expression [30, 13, 43, 12, 7, 33].

We are not overly interested in visualization [19, 2, 20, 31, 44]. We are not interested simply in querying the existing databases [6, 15]. We are not restricted to systems biology modeling pathways [4, 35, 25].

Instead, we want to exploit the fact that the object underlying a pathway diagram is a mathematical graph. Thus, we can apply algorithms from mathematics and computer science to compute invariants that describe the structure and topology of that graph, with special focus on algorithms to detect cycles. (An aside on terminology. In the graph-theoretic context, a “loop” is an edge that connects a node to itself, and a “cycle” is a path that starts at one node and traverses through one or more other nodes before returning to its starting point. Thus, the things that we call “feedback loops” are cycles and not loops in graph theory.) To implement this process, we have developed an R package called `WayFindR` that can convert the XML files from WikiPathways or KEGG into fully realized graph objects in R. These graphs can then be studied using existing R packages like `igraph` [10] or `Rgraphviz` [16] that have already implemented many graph-theoretic algorithms. There have been a few previous attempts to understand the graph-theoretic properties of biological pathways [32, 39, 28, 27]. To the best of our knowledge, however, ours is the first attempt to apply these tools systematically to better understand the presence and extent of negative feedback in biological pathways.

## 2. Methods

All computations were performed in R version 4.5.2 (2025-10-31 ucrt) of the R Statistical Software Environment [37].

### 2.1. WayFindR package framework

`WayFindR` (version 0.7.0) is a unique and robust tool. Unlike existing packages such as `rWikiPathways` [42], `pathview` [26], and `graphite` [41, 40], which focus on retrieving and visualizing pathway data, `WayFindR` emphasizes the transformation of these pathways into computationally tractable graph objects.

The `WayFindR` workflow is shown in **Figure 2**. To get started, first download a Graphical Pathway Markup Language (GPML) file from WikiPathways. (Alternatively, download an XML file from KEGG that uses their KGML format.) `WayFindR` uses the `XML` R package (version 3.99.0.19) to interpret GPML or KGML files. The core feature of `WayFindR` is its ability to convert these pathway files into the format used by R’s `igraph` package (version 2.2.1). After the conversion, researchers can apply any `igraph` function to compute graph metrics on biological pathways. In addition, `WayFindR` computes a list of cycles (if any exist) found in the directed graph. Each cycle represents a closed loop of vertices, identifying paths that begin and end at the same vertex. `WayFindR` can also interpret these cycles, allowing users to extract details such as gene labels, edge IDs, and edge types (for example, stimulatory or inhibitory). A complete, fully reproducible working example of this analysis—including the code for pathway retrieval, graph construction, cycle enumeration, feature extraction, and visualization—is provided in the `WayFindR` package vignette. These `igraph` objects can be incorporated into Cytoscape using the `createNetworkFromIgraph` function from the `RCy3` R package, which provides programmatic control of Cytoscape (import, layout, styling) from R.

**Fig. 2.**
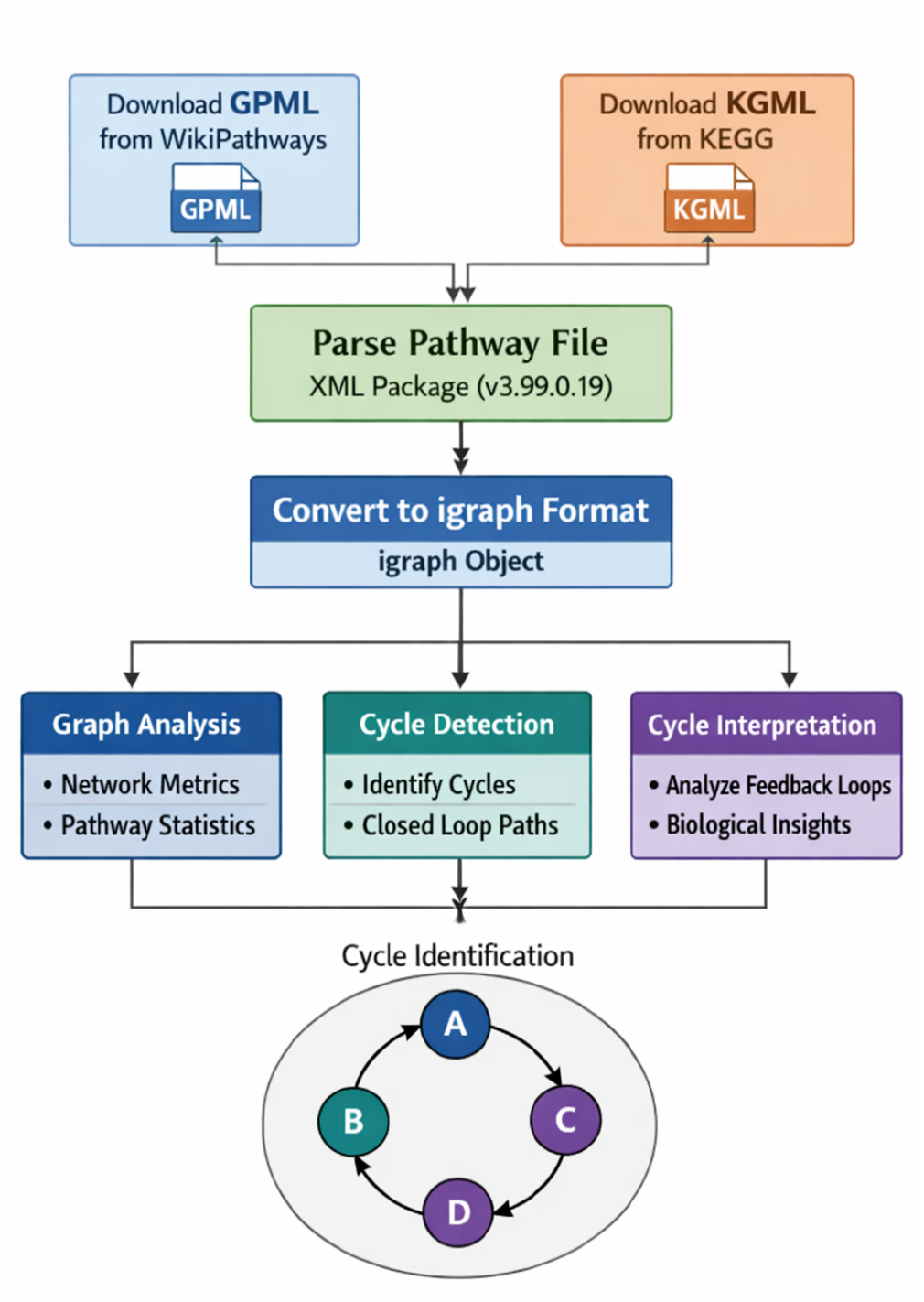
WayFindR package design and workflow.

One potential complication with the conversions is related to edge types. WikiPathways supports two standards for describing interactions between gene products: the Molecular Interaction Map (MIM) [23] and the System Biology Graphical Notation (SBGM) [24]. Perhaps because of its participatory “wiki” structure, it does not strictly enforce either standard and permits more free-form annotations as well. KEGG, meanwhile, uses its own internally defined standard that was developed before either the MIM or SBGM standards. `WayFindR` automatically converts all edge type annotations to use the MIM standard. which is the one most commonly used across WikiPathways.

### 2.2. Predicting existence of cycles using graph metrics

For exploring cycles in graphs, we focused on a set of global graph metrics already defined in the `igraph` package. Users have the flexibility to choose which metrics they want to calculate for their research purposes. Here, however, we will concentrate on a selection of metrics that are potentially interesting (**Table 1**). After using `WayFindR` to extract all cycles from biological pathways downloaded from WikiPathways and KEGG, we compiled a per-pathway table with these graph metrics. We then applied logistic regression to test whether any of these metrics had the potential to predict the presence or absence of cycles. Analyses were performed separately for each database. Logistic regression models for human, worm, mouse, and yeast were constructed using the `glm` function from the `stats` R package (version 4.5.2).

**Table 1.** Global graph metrics computed for all extracted graphs representing human, worm, mouse and yeast pathways.

| Metric | Description |
| --- | --- |
| Number of vertices | Integer count |
| Number of edges | Integer count |
| Number of negative interactions | Number of inhibitory processes/edges |
| Presence of loops | Existence of a loop edge, i.e. edge from a vertex to itself |
| Presence of multiple edges | Existence of multiple edges between same pair of vertices |
| Presence of Eulerian path or cycle | A path that passes through every edge of the graph exactly once |
| Number of clusters | Integer count |
| Density | Ratio between the number of edges and the number of theoretically possible edges |
| Radius | Minimum distance from the graph's farthest node to any vertex along the shortest path |
| Diameter | Length of the longest geodesic |
| Girth | Length of the shortest cycle in a graph |
| Global efficiency | Average of inverse distances between all pairs of vertices |
| Average path length | Average over all possible paths |
| Number of cliques | Integer count |
| Reciprocity | Proportion of mutual connections |

### 2.3. Examination of negative feedback loops in human pathways

We used the `findCycles` function from the `WayFindR` package to find, and count, the number of feedback loops in every pathway. We then interpreted these cycles using the `interpretCycle` function to extract details about the genes and interactions involved. We calculated the length of each cycle by counting the number of nodes involved. To examine the presence of nested loops within the cycles, we compared each pair of cycles within a pathway to determine if one cycle was entirely contained within another. The total number and percentage of nested loops were calculated. We also counted the number of negative (i.e., inhibitory) edges in each pathway.

We defined a “motif” as the ordered set of genes contained in a cycle, and we examined the occurrence of specific motifs across different pathways and summarized their distribution. We also calculated the frequency of each gene’s appearance in negative feedback loops, enabling us to identify the most frequently occurring genes in these regulatory cycles.

### 2.4. Gene Ontology

We used the `clusterprofiler` R package (version 4.18.4) to perform gene enrichment analysis on the set of genes involved in negative feedback loops. We performed these analyses on the Gene Ontology (GO) database, which categorizes gene properties into three main categories: Biological Process (BP), Molecular Function (MF), and Cellular Component (CC) [17]. Results of these analyses were visualized using the `enrichplot` R package (version 1.30.5).

## 3. Results

### 3.1. WikiPathways

#### 3.1.1. Example of generating graph structures from WikiPathways files

To demonstrate the basic capabilities of `WayFindR`, we retrieved the GPML file for the pathway WP3850 titled “Factors and pathways influencing insulin-like growth factor (IGF1)-Akt signaling”. We identified all cycles within the pathway and visualized the subgraph of the graph comprised of cycles. The full pathway figure and the cycle subgraph are shown in **Figure 3**.Here, the node labeled “Group1” corresponds to the pair of genes SMAD2 and SMAD3.

**Fig. 3.**
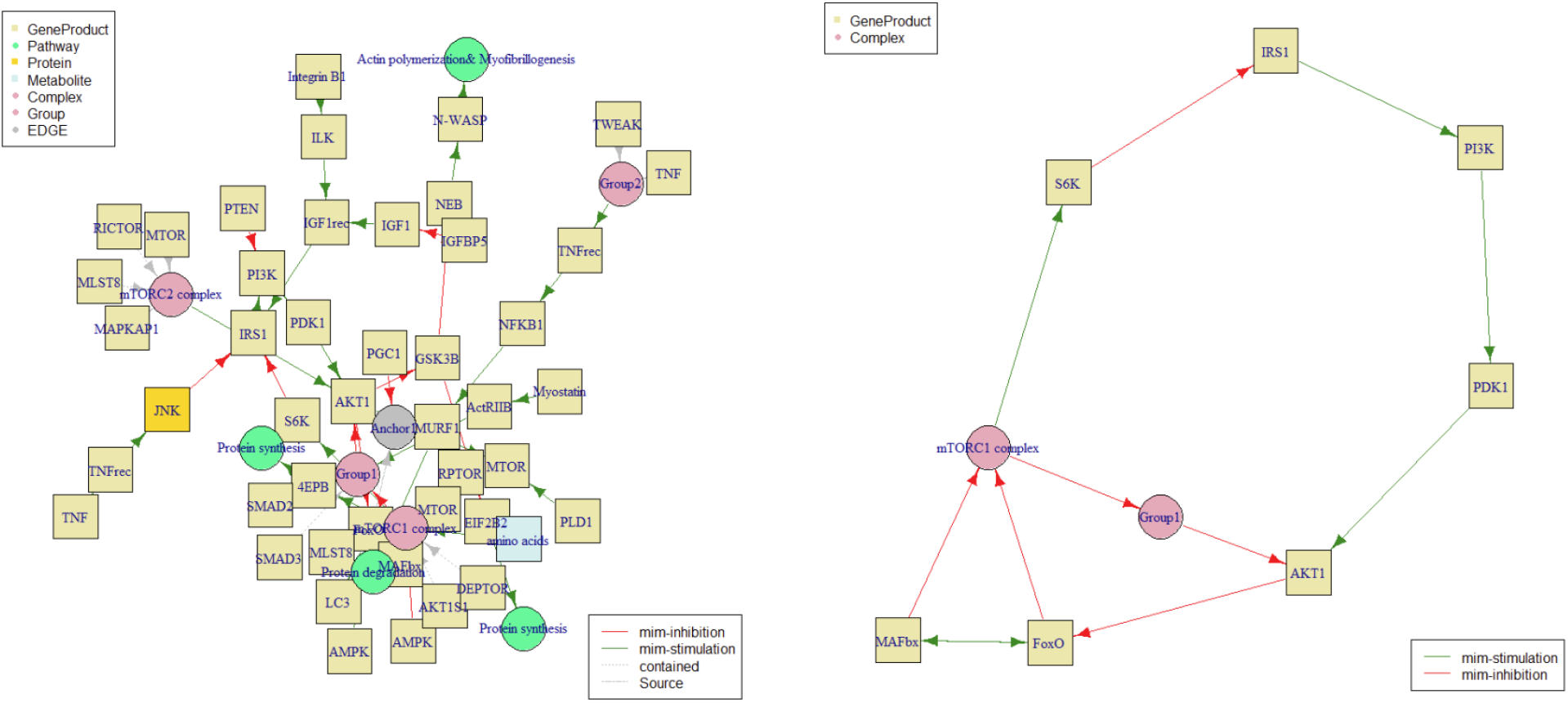
Graph (left) and cycle subgraph (right) of pathway WP3850 associated with IGF1-AKT signaling. The cycles in the chosen pathway include only inhibitory and stimulatory processes.

The pathway exhibits several cycles characterized by intricate regulatory interactions among key genes and molecular components. These cycles involve both inhibitory and stimulatory effects mediated by various genes such as PI3K, PDK1, AKT1, FoxO, MAFbx, mTORC1 complex, S6K, and IRS1. We can distinguish five cycles:

- Cycle 1: Involves a sequence of stimulatory and inhibitory actions, including PI3K and PDK1 stimulation leading to AKT1 inhibition, followed by stimulation of FoxO and mTORC1 complex, and inhibition of MAFbx, S6K, and IRS1.
- Cycle 2: Includes similar genes like PI3K, PDK1, AKT1, FoxO, mTORC1 complex, S6K, and IRS1, with variations in their regulatory interactions, highlighting different modes of stimulation and inhibition within the pathway.
- Cycle 3: Focuses on FoxO and MAFbx, illustrating specific stimulatory interactions between these genes.
- Cycle 4: Demonstrates complex interactions involving FoxO, MAFbx, mTORC1 complex, Group1, and AKT1, with a notable pattern of inhibition among these components.
- Cycle 5: Shows additional inhibitory interactions primarily involving FoxO, mTORC1 complex, Group1, and AKT1, suggesting a regulatory feedback mechanism within the pathway.

These cycles collectively depict the dynamic regulatory network within the pathway where genes interact through a combination of stimulatory and inhibitory signals to modulate cellular processes implicated in disease pathogenesis.

#### 3.1.2. Presence of negative feedback loops

We used `WayFindR` to convert all pathways for four species from the GPML format used by WikiPathways into the igraph format in R (**Table 2**). Of the 798 human (Homo sapiens, HSa) pathways, 275 (34.5%) contained cycles, 455 (57%) contained at least one inhibitory edge, and only 175 (21.9%) contained both a feedback loop and an inhibitory edge. Similar results were obtained for mouse (Mus musculus; MMu), where 21 of 169 pathways (19.5%) contained both inhibitory edges and feedback loops. In worm (Caenorhabditis elegans; CEl), only 6 out of 48 pathways contained inhibitory edges and feedback loops, while in yeast (Saccharomyces cerevisiae; SCe), only 2 out of 76 pathways (2.6%) contained inhibitory edges and feedback loops.

**Table 2.** Number of pathways with feedback loops and negative (inhibitory) edges in human (HSa), mouse (MMu), worm (CEl) and yeast (SCe) pathways.

| Species | Pathways | Cycles | Negative Edges | Negative Loops |
| --- | --- | --- | --- | --- |
| Human | 798 | 275 | 455 | 175 |
| Mouse | 169 | 56 | 90 | 33 |
| Worm | 48 | 10 | 23 | 5 |
| Yeast | 76 | 10 | 5 | 2 |

#### 3.1.3. Logistic regression

We applied logistic regression, followed by stepwise feature selection using the Akaike Information Criterion, across pathways within species to see if any graph metrics were able to predict the presence of cycles or feedback loops. The logistic regression model did not identify any metrics as significant predictors of cycles in CEl pathways. Significant predictors associated with the presence of cycles in Hsa, MMu and SCe pathways are detailed below (**Table 3**).

**Table 3.** Significant predictors for the presence of cycles in human (HS), mouse (MM), and yeast (SC) pathways. ‘P’ indicates p-value; ‘Coef’ indicates logistic regression coefficient; ‘n.s.’ means not significant.

| Predictor | Human (HS) |  | Mouse (MM) |  | Yeast (SC) |  |
| --- | --- | --- | --- | --- | --- | --- |
|  | P | Coef | P | Coef | P | Coef |
| # clusters | n.s. | n.s. | 0.02797 | 0.1976 | n.s. | n.s. |
| # negative edges | 0.01989 | -0.0532 | n.s. | n.s. | n.s. | n.s. |
| Cliques | 1.62e-05 | 1.168 | 0.02881 | 1.171 | 0.00387 | 3.624 |
| Density | 4.35e-13 | -205.5 | n.s. | n.s. | 0.00763 | -303.329 |
| Efficiency | 1.80e-14 | 90.78 | 0.02009 | 51.48 | 0.00721 | 124.855 |
| Path length | n.s. | n.s. | 0.00148 | 0.8146 | n.s. | n.s. |
| Presence of loops | 0.00845 | 0.7312 | n.s. | n.s. | n.s. | n.s. |
| Radius | 0.03528 | -0.0791 | 0.03981 | -0.2206 | n.s. | n.s. |

In general, the coefficients in logistic regression indicate the change in the log odds of the outcome variable for a one-unit increase in the corresponding predictor variable. A positive coefficient suggests that the predictor increases the likelihood of the outcome variable falling into a specific category, whereas a negative coefficient suggests it decreases the likelihood [34].

In our study, “efficiency” and “cliques” appear as significant predictors in all three species. The findings indicate that human pathways with more negative edges are less likely to exhibit cycles. Additionally, pathways with loops (self-edges) show a strong association with the presence of cycles. A higher density, which reflects how interconnected the network is, correlates significantly with a decrease in the likelihood of cycles, with a substantial negative coefficient of 205.5. This suggests that denser networks are markedly less likely to have cycles. Pathways with larger radii are also less likely to exhibit cycles. Conversely, more efficient networks are significantly more likely to contain cycles, and pathways with more cliques exhibit a higher likelihood of cycles. In mouse pathways, more clusters and longer average path lengths increase the likelihood of cycles occurring. Smaller radius and higher efficiency also show a positive correlation with cycle presence. Pathways with more cliques tend to exhibit cycles. Similarly, in yeast pathways, significant predictors for cycle presence include lower density, higher efficiency, and more cliques.

#### 3.1.4. Properties of Negative Feedback Loops

Next, our focus shifted to negative feedback loops in human pathways from the WikiPathways database. Out of 275 human pathways that contained any type of cycle, only 82 had inhibitory negative-feedback edges. We found that the length of negative feedback loops ranges from a minimum of 2 nodes to a maximum of 50. Specifically, in the case of WP4718, which relates to ‘Cholesterol metabolism with Bloch and Kandutsch-Russell pathways,’ we found a negative feedback loop containing 50 nodes. Overall, nearly 40% of the nodes in negative feedback loops correspond to gene products, while approximately 3% belong to an undefined category. The top three types of edges observed in negative feedback loops are “mim-inhibition”, “mim-stimulation”, and “none”. Summary of these observations is depicted in **Figure 4**.

**Fig. 4.**
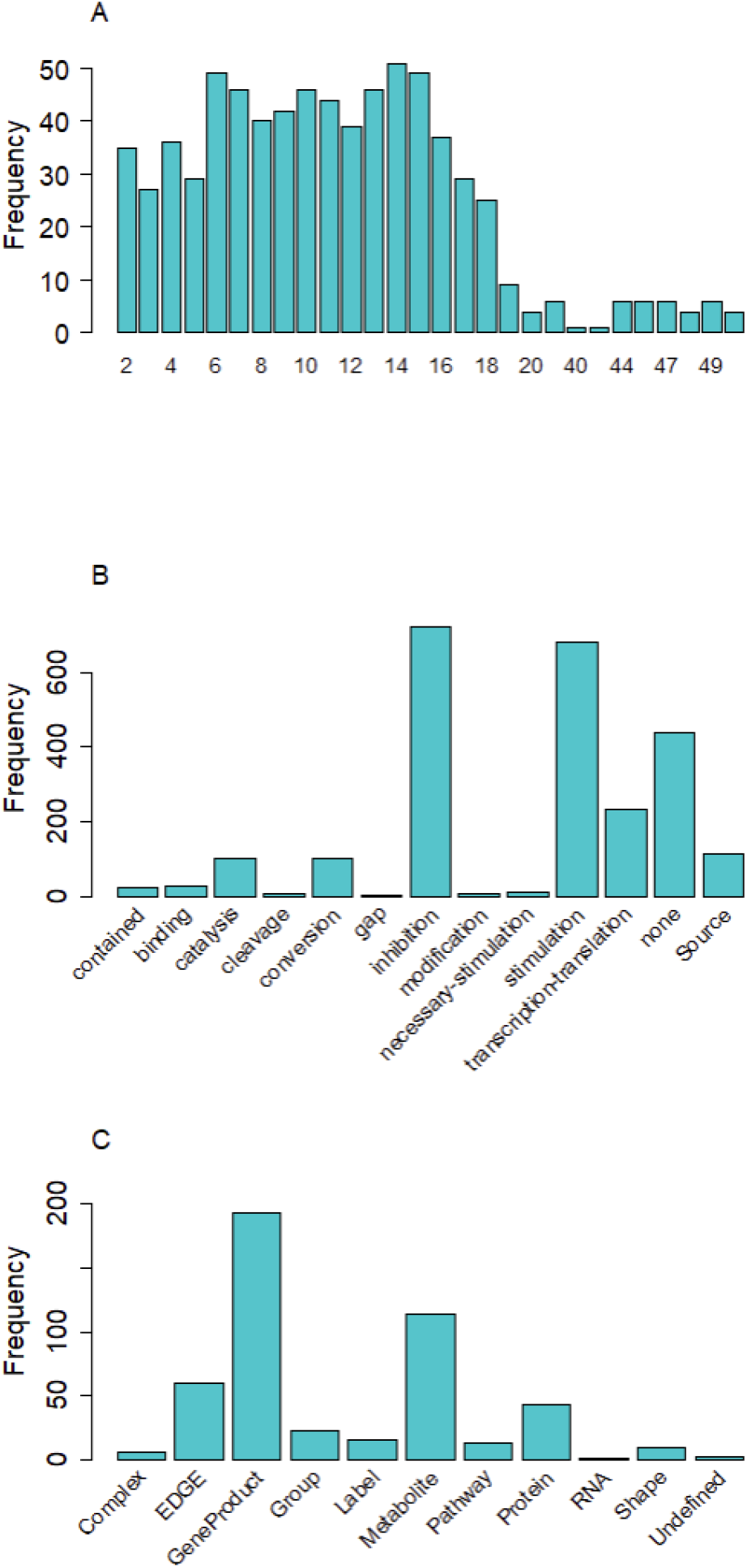
(A) Length of negative feedback loops. (B) Distribution of edges types. (C) Distribution of node types in negative feedback loops. A detailed description of WikiPathways GPML terminology is available in the CRAN package vignette and the official WikiPathways GPML vocabulary (https://vocabularies.wikipathways.org/gpml).

In our analysis of nested negative feedback loops, we evaluated the structure of cycles within each pathway graph to identify loops that are nested within other loops. Across all pathway graphs with negative feedback loops analyzed, the total number of loops identified was 722. Out of these, 489 (67.73%) loops were found to be nested within other loops. The high percentage of nested loops indicates a significant level of hierarchical organization within the negative feedback loops of these pathways.

#### 3.1.5. Note on WP4718: “Cholesterol Metabolism with Bloch and Kandutsch-Russell Pathways”

Our analysis initially identified a 50-node, representation-driven cycle tracing the linear cholesterol biosynthesis cascade from acetoacetyl-CoA to cholesterol. Most edges in this cycle correspond to biochemical conversions (mim-conversion), interleaved with Anchor elements and Source import labels— schema-level, non-regulatory features by design. Within this topology, a compact regulatory core is present: 25-hydroxycholesterol inhibits SREBF2 (mim-inhibition), and SREBF2 stimulates multiple biosynthetic steps (mim-stimulation), with catalysis links (mim-catalysis) indicating enzyme–reaction associations. By applying our pre-specified valid-loop criteria (molecular nodes + signed regulatory edges only; excluding conversion, Anchor/Source, and other non-regulatory link types), the long, graph-theoretic cycle is refined to a short, biologically meaningful negative-feedback motif: sterols *→* | SREBF2 *→* biosynthetic enzymes. This example illustrates that the apparent large cycle reflects curated pathway structure rather than regulatory complexity, while `WayFindR` faithfully preserves the input topology and provides filters that allow users to focus on true regulatory interactions.

#### 3.1.6. Motifs

The analysis of motifs in negative feedback loops reveals the presence of key regulatory interactions that are conserved across multiple pathways. The motif “MDM2-TP53” is the most prevalent, found in four pathways: WP1742, WP2261, WP2446, and WP5087. This motif is of particular interest as MDM2 is known to inhibit TP53, a critical regulatory relationship in cellular pathways [18]. The motif “IRF6-TP63” was identified in the pathways WP3655 and WP4674, suggesting another key regulatory interaction.

#### 3.1.7 Genes frequently involved in feedback loops

Next, we investigated which genes appear most frequently in negative feedback loops across pathways. The gene TP53 was identified as prominently participating in negative feedback loops across various pathways, specifically in TP53 network (WP1742), Glioblastoma signaling (WP2261), Retinoblastoma gene in cancer (WP2446), MicroRNA network associated with chronic lymphocytic leukemia (WP4399), NAD metabolism in oncogene-induced senescence and mitochondrial dysfunction-associated senescence (WP5046), TCA cycle in senescence (WP5050), and Pleural mesothelioma (WP5087). The biological roles and implications of TP53 are well-documented. It acts as a tumor suppressor in many types of tumors, inducing growth arrest or apoptosis depending on the physiological context and cell type. TP53 is involved in cell cycle regulation by negatively regulating cell division through trans-activation of a set of genes necessary for this process [18]. Mutations in TP53 are associated with various human cancers, including hereditary cancers such as Li-Fraumeni syndrome.

#### 3.1.8. Gene enrichment analysis

We performed gene enrichment analysis to identify overrepresented Gene Ontology (GO) terms among the genes that appeared in at least one negative feedback loop. This analysis highlighted significant biological processes and molecular functions associated with this gene set. The enriched terms were visualized using the `cnetplot` function from the `enrichplot` R package. (**Figure 5**).

**Fig. 5.**
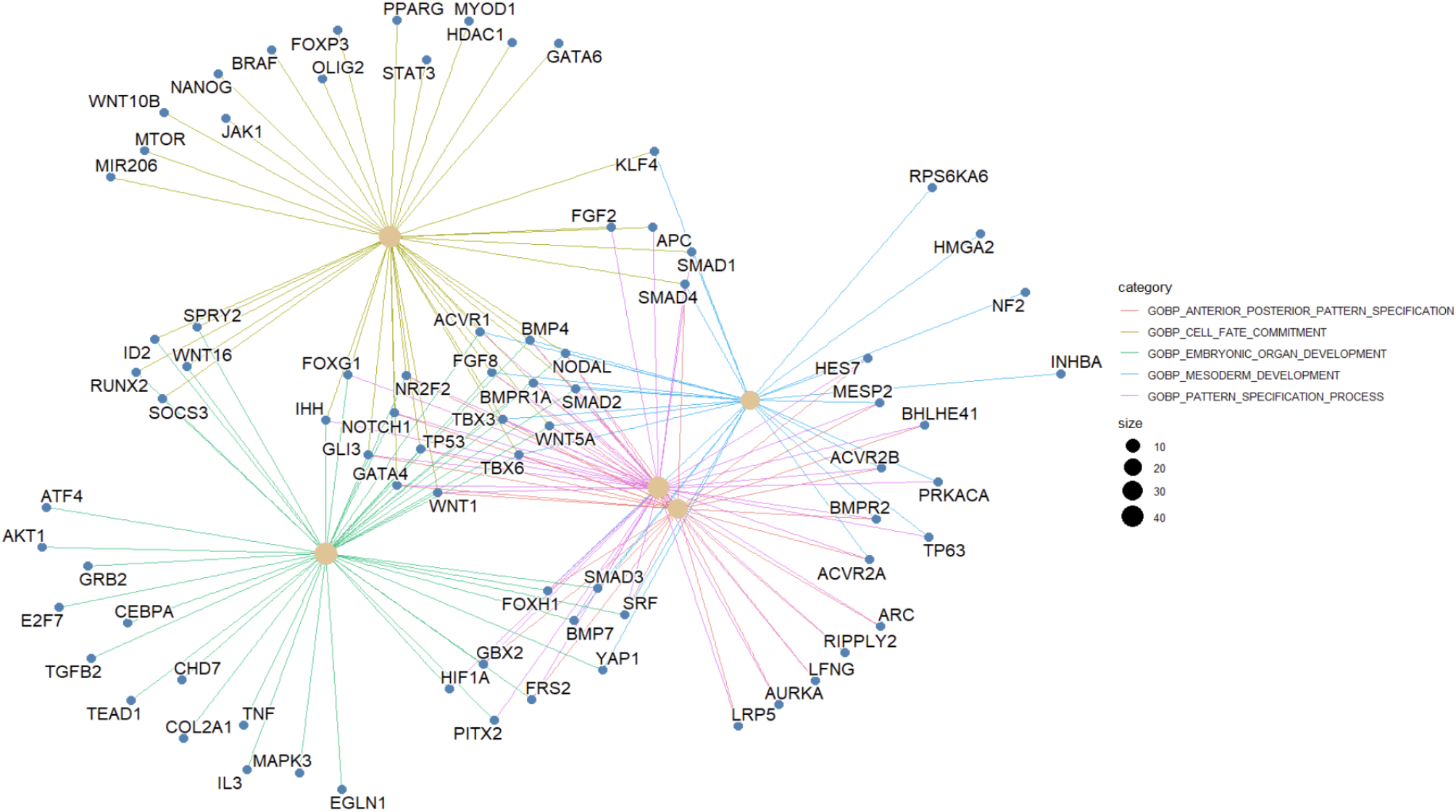
The linkages of overrepresented genes in negative feedback loops and biological concepts (GO terms) as a network.

**Fig. 6.**
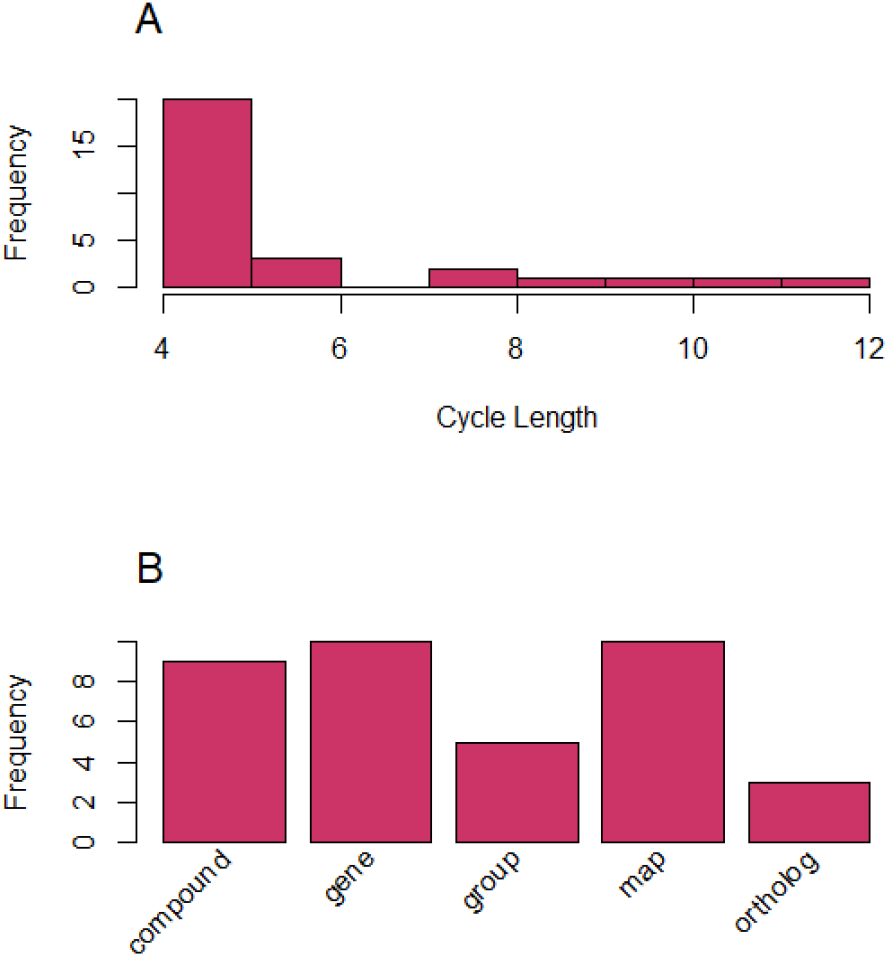
(A) Distribution of negative feedback loop lengths; (B) Frequency of node types involved in negative feedback loops across KEGG human pathways.

**Fig. 7.**
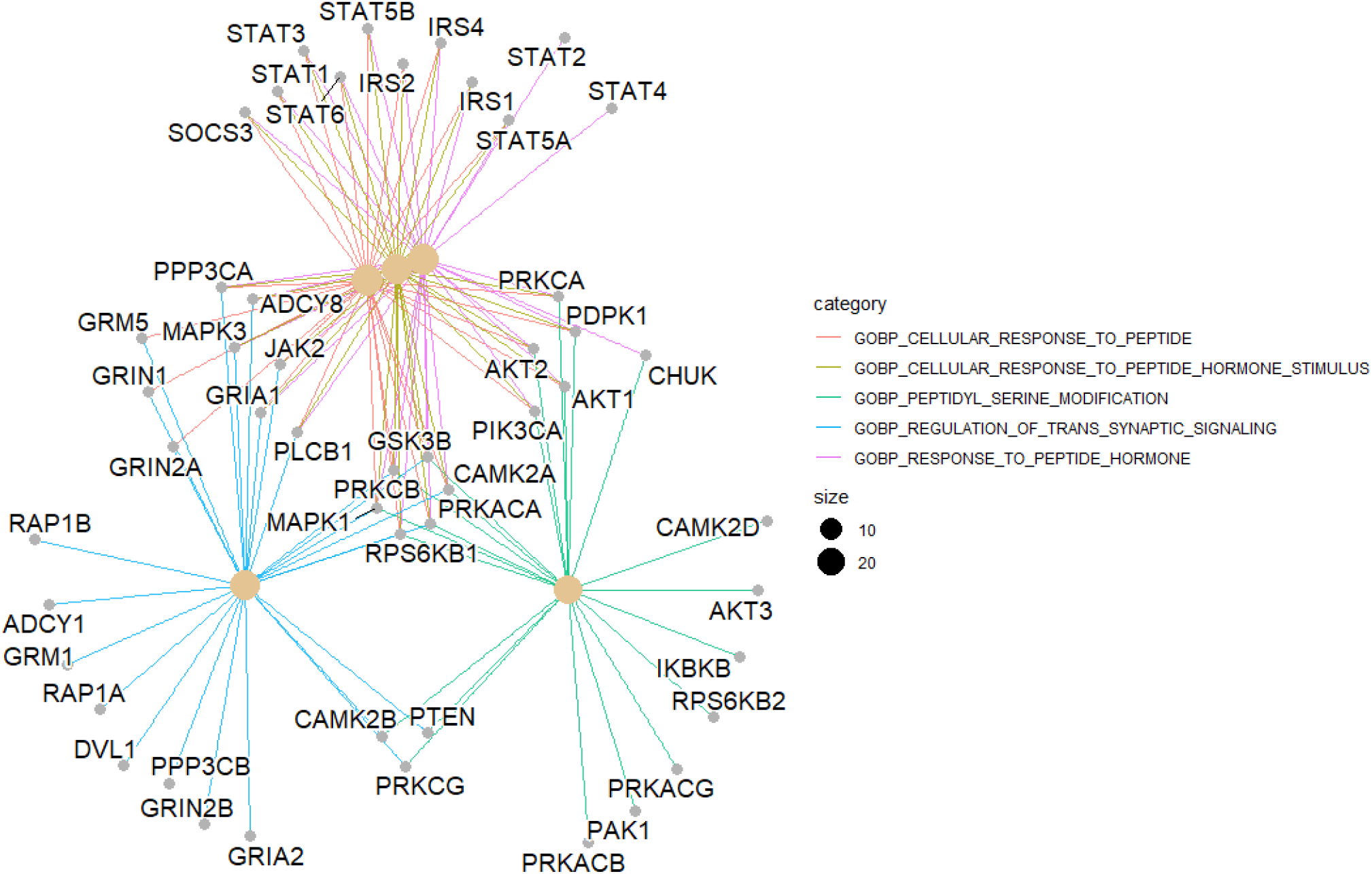
The linkages of overrepresented genes in negative feedback loops extarcted from KEGG human pathways and biological concepts (GO terms) as a network.

Interestingly, all overrepresented terms came from the Gene Ontology Biological Process (GOBP) category, with no significant enrichment of Molecular Function or Cellular Process categories. Two of the five GOBP terms pertain to developmental processes: mesoderm development and embryonic organ development. The remaining terms are related to anterior-posterior pattern specification, cell fate commitment, and pattern specification processes. Among these, mesoderm development and anterior-posterior pattern specification have fewer associated genes. Eight genes — BMP4, BMPR1A, FGF8, NODAL, SMAD2, TBX3, TBX6, and WNT5A — are present across all five GOBP categories. TP53, the gene most frequently found in negative feedback loops, is associated with four annotation categories.

### 3.2. KEGG

While a detailed analysis of feedback loops was conducted on WikiPathways, a comparable analysis of 328 KEGG human pathways revealed similar trends. Only 10 out of 91 pathways with cycles contained inhibitory interactions forming negative feedback loops, confirming the rarity of such regulatory motifs. A full summary of the KEGG analysis is provided in the Supplementary Materials.

## 4. Discussion and conclusion

In this paper, we introduce a novel package called “WayFindR” designed to address challenges in analyzing biological pathways, with a particular focus on feedback loops.

For a given pathway, `WayFindR` can identify and analyze cycles within the pathway graph, offering insights into the interconnections and feedback mechanisms present. By interpreting these cycles within the associated graph context, `WayFindR` helps researchers understand how cyclic structures contribute to the overall dynamics and regulation of the pathway. Additionally, `WayFindR` provides a comprehensive suite of functions to calculate various graph metrics. These metrics are crucial for evaluating the structural and functional properties of biological pathways, facilitating a deeper understanding of how individual components interact and influence the pathway’s behavior.

Despite the extensive number of pathways studied across various species, a surprisingly small proportion exhibit feedback loops. In WikiPathways, only 34.46% (275/798) of Homo sapiens pathways contain cycles; this fraction is 33.73% (57/169) in Mus musculus, 20.83% (10/48) in Caenorhabditis elegans, and 13.16% (10/76) in Saccharomyces cerevisiae. A parallel KEGG analysis shows 27.7% (91/328) of human pathways contain cycles, with 33.1% (56/169) in mouse, 20.8% (10/48) in worm, and 13.2% (10/76) in yeast; pathways with at least one negative feedback loop are 3.0% (10/328) in human, 19.5% (33/169) in mouse, 10.4% (5/48) in worm, and 2.6% (2/76) in yeast (Table 4).

**Table 4.** Overview of the KEGG database: Number of pathways with feedback loops and negative (inhibitory) edges in human (HSa), mouse (MMu), worm (CEl) and yeast (SCe) pathways.

| Species | Pathways | Cycles | Negative Edges | Negative Loops |
| --- | --- | --- | --- | --- |
| Human | 328 | 91 | 167 | 10 |
| Mouse | 169 | 56 | 90 | 33 |
| Worm | 48 | 10 | 23 | 5 |
| Yeast | 76 | 10 | 5 | 2 |

**Table 5.**
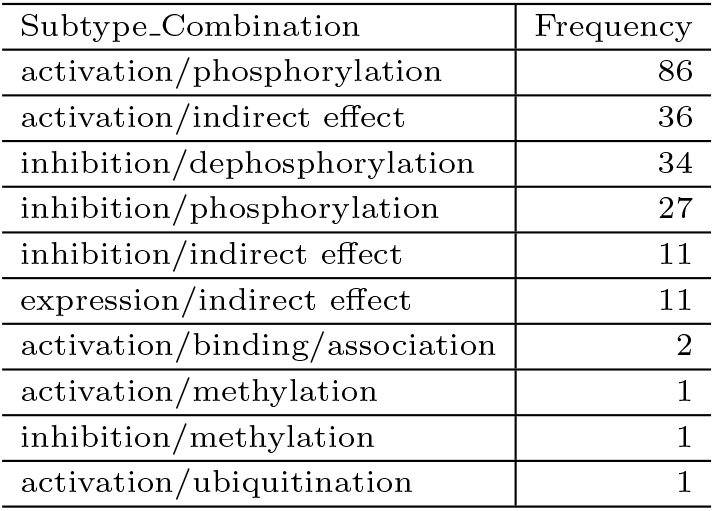
Subtype pair frequencies in KEGG human pathways exhibiting negative feedback.

The relatively limited number of identified negative feedback loops likely arises from two main factors. First, negative feedback loops remain underexplored due to historical research focus and methodological limitations, resulting in incomplete characterization within existing pathway databases. These findings highlight the rarity or potentially underexplored nature of feedback loops in biological pathways across different organisms. This rarity may reflect either the inherent complexity and specificity of these loops, making them challenging to identify and characterize, or a gap in current research methodologies and knowledge. Further studies are needed to uncover and understand the dynamics of feedback loops in biological systems. Second, technical challenges associated with databases like KEGG and WikiPathways may contribute to underreporting. Many pathways do not fully comply with annotation standards or lack comprehensive, standardized representations of interactions—particularly inhibitory or negative regulatory edges essential for detecting feedback loops. These inconsistencies and gaps in pathway curation impede accurate identification and analysis, highlighting the need for improved data quality and standardized annotation to better understand these critical regulatory motifs.

Our logistic regression analysis identified significant predictors for the presence of cycles in human, mouse, and yeast pathways sourced from the WikiPathways repository. Notably, network efficiency and the number of cliques emerged as consistent predictors across all species, suggesting common structural features associated with feedback loops. The presence of negative feedback loops in a substantial number of pathways underscores their importance in maintaining cellular homeostasis and regulating complex biological processes.

A detailed examination of negative feedback loops in human pathways revealed diverse structural characteristics and interaction patterns. We found that the “MDM2-TP53” motif is the most frequently observed in these loops. Recognizing that MDM2-TP53 is a common motif in feedback loops could provide insights into targeting these interactions for therapeutic purposes, particularly in cancer research where TP53 is often mutated or dysregulated.

Our gene set enrichment analysis offers valuable insights into the functional roles of genes involved in negative feedback loops and their impact on critical biological processes. The connection between these genes and developmental as well as pattern specification processes highlights the intricate relationship between regulatory mechanisms and cellular functions. Differences in the gene universe represented by WikiPathways and KEGG, plus schema-level choices that affect which genes participate in loops, naturally lead to different enriched GO terms when analyses are run per resource; we present these enrichment results as exploratory and include full term lists and adjusted p-values alongside figures for transparency.

Overall, results from both WikiPathways and KEGG support the conclusion that negative feedback loops are present in only a small fraction of pathways, suggesting their selective deployment in tightly regulated systems.

In conclusion, studying feedback loops, especially negative ones, provides profound insights into the regulatory mechanisms governing biological systems. Tools like WayFindR enhance our capability to explore these complex structures, paving the way for new discoveries and innovative therapeutic approaches. Integrating biological knowledge with computational tools offers a robust framework for advancing our understanding of complex biological networks and their roles in health and disease.

## 5. Supplementary Materials

### 5.1. KEGG

We extracted 328 human pathways from the KEGG database and systematically analyzed them for the presence of feedback loops. Among these, 91 pathways contained at least one cyclic structure. Of those, only 10 pathways featured cycles with at least one inhibitory edge. In total, we identified 23 negative feedback loops across these 10 pathways.

The lengths of negative feedback loops ranged from 4 to 12 nodes. The most common node types within these loops were “gene” and “map”’ with “activation/phosphorylation” being the predominant edge subtypes connecting them.

Moreover, we checked how many times each gene appeared across all 23 feedback loops. This revealed that 97 unique genes are involved, with CISH and STAT3 being the most frequently recurring, each appearing 10 times. This high recurrence aligns with their known roles in cytokine signaling pathways. Specifically, STAT3 is activated by cytokines and translocates to the nucleus to induce target genes, including CISH. In turn, CISH acts as a feedback inhibitor by suppressing further STAT signaling.

Functional enrichment analysis of the 97 genes participating in negative feedback loops revealed a significant overrepresentation of cellular and hormonal response processes. The most enriched categories included cellular response to peptide and response to peptide hormone. Additionally, enrichment in regulation of trans-synaptic signaling and peptidyl-serine modification indicates a role in neuronal communication and post-translational modifications, both of which are critical for fine-tuning dynamic responses in complex signaling systems.

Comparison with the DrugBank database revealed 45 genes involved in negative feedback loops that overlap with known drug targets.

## 7. Acknowledgements

## Notes

### Competing Interest Statement

The authors have declared no competing interest.

### Funding

- Georgia Cancer Center at Augusta University: Starup funds for KRC

